# Cerebellar Neuronal Dysfunction Accompanies Early Motor Symptoms in Spinocerebellar Ataxia Type 3

**DOI:** 10.1101/2021.04.29.441865

**Authors:** Kristin Mayoral-Palarz, Andreia Neves-Carvalho, Patrícia Maciel, Kamran Khodakhah

## Abstract

Spinocerebellar Ataxia Type 3 (SCA3) is an adult-onset, progressive ataxia. SCA3 presents with ataxia before any gross neuropathology. A feature of many cerebellar ataxias is aberrant cerebellar output that contributes to motor dysfunction. We examined whether abnormal cerebellar output was present in the CMVMJD135 SCA3 mouse model and, if so, whether it correlated with the disease onset and progression. *In vivo* recordings showed that the activity of deep cerebellar nuclei neurons, the main output of the cerebellum, was altered. The aberrant activity correlated with the onset of ataxia. However, although the severity of ataxia increased with age, the severity of the aberrant cerebellar output was not progressive. The abnormal cerebellar output, however, was accompanied with non-progressive abnormal activity of their upstream synaptic inputs, the Purkinje cells. *In vitro* recordings indicated that alterations in both intrinsic Purkinje cell pacemaking and in their synaptic inputs contributed to abnormal Purkinje cell activity. These findings implicate abnormal cerebellar physiology as an early, consistent contributor to pathophysiology in SCA3, and suggest that the aberrant cerebellar output could be an appropriate therapeutic target in SCA3.

**Summary Statement:** In a mouse model of Spinocerebellar ataxia type 3 aberrant cerebellar physiology is apparent early in disease, before overt neuronal pathology or neuronal death.

## Introduction

The cerebellum is involved in motor coordination and maintenance of balance. Dysfunction of the cerebellum can lead to ataxia, or uncoordinated movement. The most common dominantly inherited ataxia is Spinocerebellar Ataxia Type 3 (SCA3), also known as Machado-Joseph Disease. SCA3 is caused by a heterozygous (CAG)n/polyglutamine expansion of the Ataxin-3 gene (Kawaguchi *et al*., 1994). Patients present with adult-onset progressive ataxia, along with a combination of ophthalmoplegia, spasticity, dystonia, muscular atrophy, or other extrapyramidal signs (Coutinho & Andrade, 1978; Barbeau *et al*., 1984). The most common affected regions on pathology include the cerebellar dentate nucleus, pallidum, substantia nigra, thalamus, subthalamic nuclei, red nuclei, and, to a lesser extent, the cerebellar cortex (Woods & Schaumburg, 1972; Rosenberg *et al*., 1976; Romanul *et al*., 1977; Stefanescu *et al*., 2015; Hernandez-Castillo *et al*., 2018; Koeppen, 2018; Wang *et al*., 2020). Degeneration and other pathologic hallmarks, such as protein aggregate formation, occur late in the course of the disease.

In the cerebellum of SCA3 patients, pathology has consistently shown late-onset degeneration of the deep cerebellar nuclei (DCN), the main output of the cerebellum, but little to no degeneration of the Purkinje cells, the main computational unit of the cerebellar cortex (Rosenberg *et al*., 1976; Haines & Dietrichs, 2012; Koeppen, 2018). It is plausible that cerebellar neuronal dysfunction contributes to the early motor phenotype. The cerebellum encodes movement related information through rapid increases or decreases in the firing rate of the output nuclei, the DCN (Thach, 1968; Eccles, 1973). The DCN neurons are intrinsically active (Raman *et al*., 2000), meaning they fire in the absence of synaptic input, and fire with a regular firing pattern (Alvina & Khodakhah, 2008). If the precision of spontaneous activity is altered, the accuracy of encoding synaptic information will be impaired. The DCN receive synaptic input from mossy fibers, climbing fibers, and Purkinje cells. Purkinje cells are also intrinsically active neurons (Raman & Bean, 1999), and have a regular firing pattern (Hausser & Clark, 1997). Disruption to the regularity of firing of Purkinje cells and the neurons within the deep cerebellar nuclei has been found in other mouse models of ataxia (Walter *et al*., 2006; Alvina & Khodakhah, 2010b; a; Tara *et al*., 2018). As Purkinje cells are the main computational unit of the cerebellar cortex and converge onto DCN, disruption to their activity can impact the ability of the output of the cerebellum to accurately encode motor-related events. This raises two questions: 1) is cerebellar neuronal activity altered in SCA3? And 2) if so, is the dysfunction progressive, and does it correlate with the motor phenotype?

A SCA3 mouse model (CMVMJD135) expresses the expanded disease-relevant ataxin-3c isoform under the human cytomegalovirus (CMV) promoter in a heterozygous manner, conferring wide expression at near endogenous levels throughout the nervous system, similar to what is seen in patients (Silva-Fernandes *et al*., 2014; Teixeira-Castro *et al*., 2015). Overall, the mouse model is ideal to study the cerebellar contribution of disease as it recapitulates the adult onset, progressive motor phenotypes seen in SCA3 patients, neuropathological findings, in addition to a late in disease neuronal degeneration profile with sparing of Purkinje cells.

In the CMVMJD135 mouse model (referred to as SCA3 mice throughout this paper) the DCN had irregular firing in awake, head restraint mice. This indicated early, non-progressive, cerebellar neuronal alteration in SCA3 from either DCN neurons, or from irregular activity upstream of the nuclei. *In vivo* Purkinje cell activity, which converges onto the DCN, were also irregular, but not progressive, likely contributing to the abnormal cerebellar output. *In vitro* acute slice recordings allowed investigation of the contributors to Purkinje cell irregular firing. Purkinje cell irregularity had both aberrant intrinsic firing and altered synaptic activity, with a significant synaptic contribution. Altogether we find that the cerebellum is a site of early dysfunction, before pathology is detected, suggesting that Purkinje cell irregular activity may be an early driver of pathophysiology and a potential therapeutic target in SCA3.

## Results

### SCA3 Mice Had Motor Dysfunction Early in Disease

To further characterize the motor phenotype of the SCA3 mice across the course of the disease, mice were assessed using established tests of motor performance not previously applied to this particular disease model at an early, mid, and late disease state. The littermate wild-types, referred to as wild-type throughout this paper, and SCA3 mice were tested on a disability scale for gross motor abnormalities (Weisz *et al*., 2005; Tara *et al*., 2018). Using this scale, a mouse with no motor symptoms receives a score of 0, a 1 indicates abnormal gait, and the severity of motor impairment scales up to a score of 5 (Figure 1A). Based on this assessment, SCA3 mice had motor impairment at 12-weeks of age (wild-type 0.17±0.14, SCA3 0.52±0.28, p=0.0018), with overt impairment at 34-weeks (wild-type 0.19±0.17, SCA3 1.70±0.58, p<0.0001) and severe impairment by 60-weeks (wild-type 0.27±0.16, SCA3 2.63±0.10, p=0.0003) (Figure 1B). This demonstrates the progressive nature of the disease in SCA3 mice. By 60-weeks, the motor symptoms are a combination of deficits from the central and peripheral nervous systems (Duarte-Silva *et al*., 2014; Silva-Fernandes *et al*., 2014; Teixeira-Castro *et al*., 2015; Esteves *et al*., 2019).

**Figure 1:**
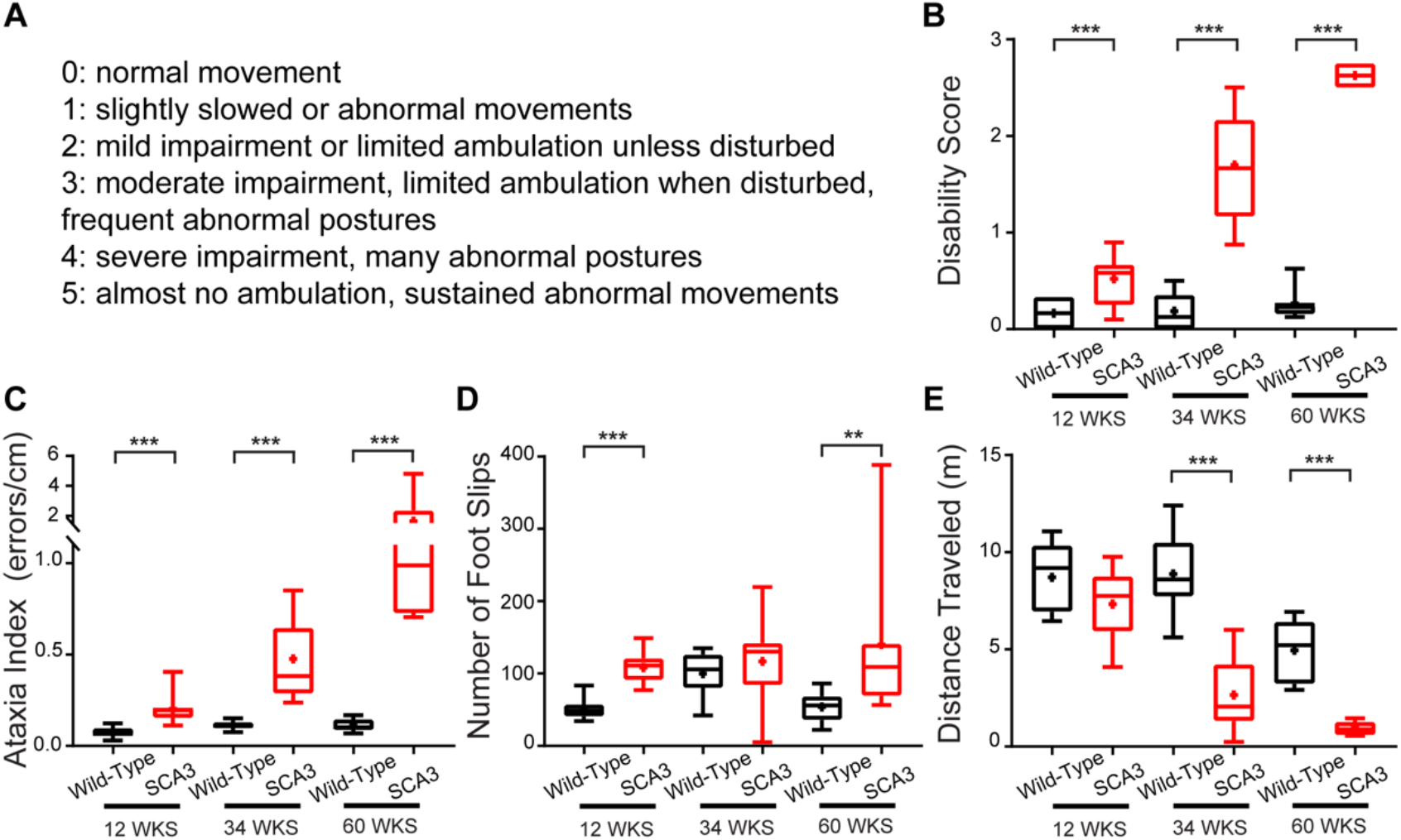
SCA3 Motor Symptoms Progress With Age, With Onset As Early As 12-weeks. SCA3 mice and wild-types were tested at 12-, 34-, and 60-weeks of age on the disability score (A-B) and parallel rod floor test (C-E). SCA3 mice have an increased disability score (A-B) compared to wild-types at 12-weeks, and the score progresses with age. SCA3 mice have an increased ataxia index (C) at 12-weeks due to increased foot slips (D) with no change in distance traveled (E). At 34-and 60-weeks of age, the increased ataxia index reflects combined decrease in distance traveled and increased foot slips. Parallel Rod Floor Test: 12-weeks: wild-type N = 9, SCA3 N = 9. 34-weeks: wild-type N = 8, SCA3 N = 9. 60-weeks: wild-type N = 8, SCA3 N = 7. Disability Score (all scores are average of 2 trials): 12-weeks: wild-type N = 10, SCA3 N = 15. 34-weeks: wild-type N = 17, SCA3 N = 19. 60-weeks: wild-type N = 8, SCA3 N = 7. **p<0.01, ***p<0.001. T-test (B – 12-weeks. D – 12-weeks, 34-weeks. E – all), Mann-Whitney test (B – 34-weeks, 60-weeks. C – all. D – 60-weeks).

The second assay, the parallel rod floor test, is more sensitive to motor incoordination and eliminates variability due to human error in assessing the disability score (Kamens *et al*., 2005; Kamens & Crabbe, 2007). The ataxia index (Figure 1C), calculated by number of foot slips (Figure 1D) per distance traveled (cm) (Figure 1E), was averaged per mouse over two trials. SCA3 mice had a higher ataxia index than wild-types as early as 12-weeks (wild-type 0.074±0.029, SCA3 0.202±0.083, p<0.0001), with the ataxia index further increasing at 34-weeks (wild-type 0.112±0.023, SCA3 0.476±0.211, p<0.0001) and 60-weeks of age (wild-type 0.111±0.035, SCA3 1.645±1.51, p=0.0003) (Figure 1C-E). Interestingly, at 12-weeks, the increased ataxia index was driven from an increased number of foot slips with no significant decrease in distance travelled, indicating motor incoordination and not an overall change in gross movement as a main contributor to the impairment (Figure D-E). While the progressive increase in ataxia index was likely due to a combination of increased central and peripheral dysfunction, the parallel rod floor test provides a sensitive measure of motor impairment in SCA3 mice as early as 12-weeks of age, an age at which no brain pathology has been detected (Silva-Fernandes *et al*., 2014; Teixeira-Castro *et al*., 2015; Esteves *et al*., 2019).

### Activity Of Deep Cerebellar Nuclei Was Altered, But Not Progressive, In SCA3 Mice

We first examined whether there are any alterations in the activity of the cerebellar output nuclei *in vivo*. To minimize the time-variant information encoded by synaptic activity, we assessed the firing of the neurons in the deep cerebellar nuclei (DCN) in stationary animals. While the cerebellar contribution to actions such as standing remain, the need for motor signals driving rapid changes in firing are reduced, allowing for a more accurate assessment of the precision of firing. Therefore, *in vivo* recordings of the DCN neurons were made from awake, head-restrained, stationary mice. The inter-spike interval coefficient of variation (ISI CV) was used to measure the regularity of firing (Hausser & Clark, 1997; Walter *et al*., 2006; Tara *et al*., 2018). The ISI CV is the standard deviation of the inter-spike interval divided by the average inter-spike interval. The larger the ISI CV, the more irregular the firing and less precise the activity.

In SCA3 mice, the ISI CV of DCN was increased, but not progressive, at 12-, 34-, and 60-weeks of age (12-weeks: wild-type 0.371±0.114, SCA3 0.648±0.199, p<0.0001; 34-weeks: wild-type 0.427±0.054, SCA3 0.599±0.170, p<0.0001; 60-weeks: wild-type 0.421±0.121, SCA3 0.599±0.152, p<0.0001) (Figure 2E). DCN neurons had a decreased average firing rate (the reciprocal of the average inter-spike interval) at 12-, 34-, and 60-weeks of age in SCA3 mice (12-weeks: wild-type 71.9±18.9, SCA3 53.4±21.5, p=0.0002; 34-weeks: wild-type 74.1±19.8, SCA3 64.7±19.8, p=0.0359; 60-weeks: wild-type 80.3±26.0, SCA3 63.9±19.2, p=0.0154) (Figure 2F). The decrease in average firing rate would be expected to result in a decreased ISI CV if the firing pattern had not changed. Therefore, with a decrease in firing rate, the increased irregularity, as reported by ISI CV, in SCA3 mice was greater than observed. Interestingly, while the ISI CV was increased at all timepoints, there was no progressive increase the ISI CV, suggesting extra-cerebellar contributions to the progressive phenotype of SCA3 mice.

**Figure 2:**
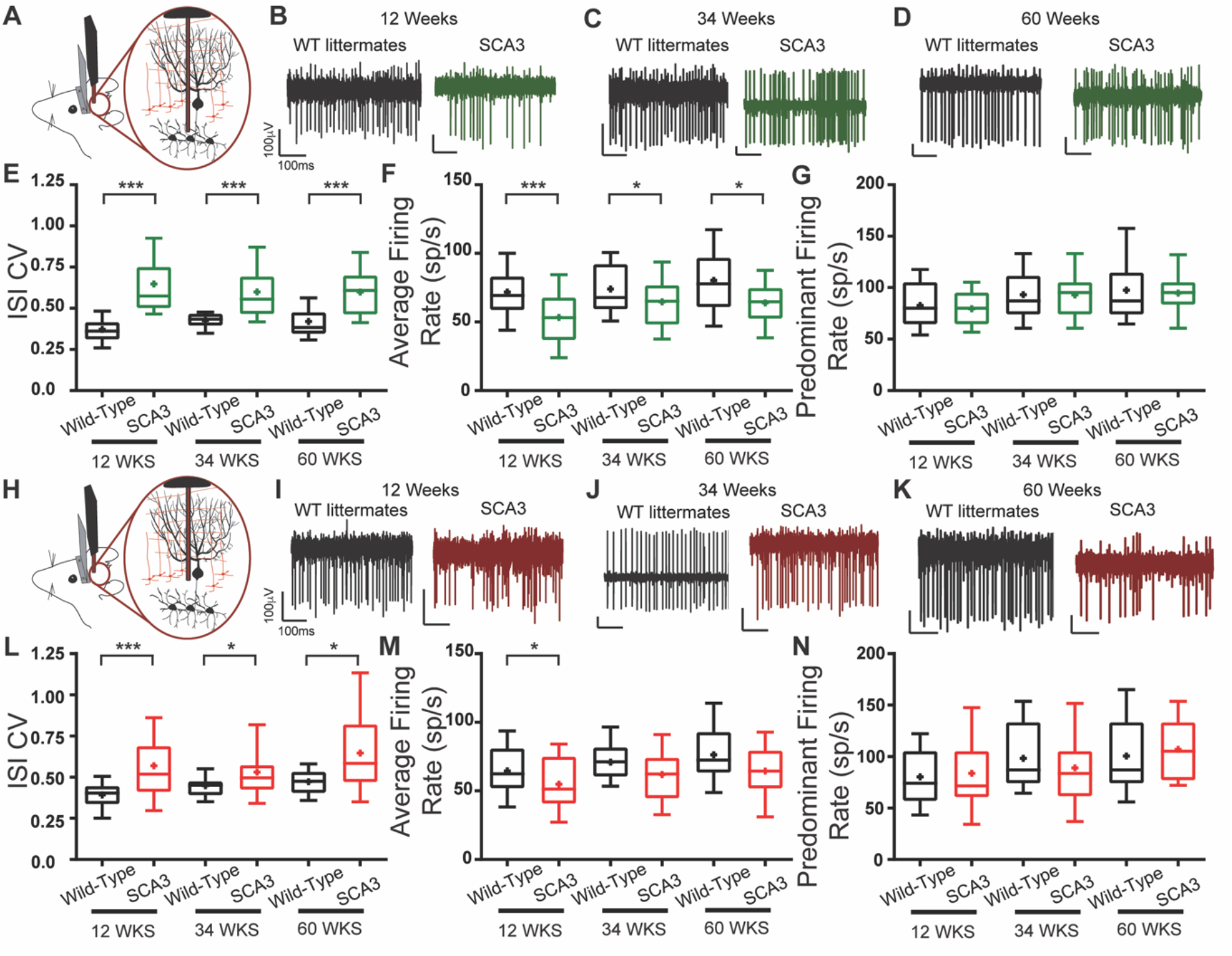
Cerebellar Neuronal Dysfunction in SCA3 Mice Across Disease Progression. *In Vivo* awake, head-fixed cerebellar deep cerebellar nuclei (A) and Purkinje cell (H) single-unit recordings were performed from SCA3 mice and wild-types at 12-, 34-, and 60-weeks of age. Example deep cerebellar nuclei firing traces from wild-type (black) and SCA3 (green) at 12-(B), 34-(C), and 60-(D) weeks of age. Deep cerebellar nuclei inter-spike interval coefficient of variation (ISI CV) (E) is increased in SCA3 while the average firing rate (F) is decreased, with no change in predominant firing rate (G) for all ages tested. Example Purkinje cell firing traces from wild-type (black) and SCA3 (red) at 12-(I), 34-(J), and 60-(K) weeks of age. Purkinje cells ISI CV (L) is increased in SCA3 while the average firing rate (M) is decreased or trends to decrease, with no change in predominant firing rate (N). Deep Cerebellar Nuclei: 12-weeks: wild-type N = 6, n = 39, SCA3 N = 8, n = 33. 34-weeks: wild-type N = 12, n = 33, SCA3 N = 16, n = 65. 60-weeks: wild-type N = 5, n = 24, SCA3 N = 3, n = 25. Purkinje cells: 12-weeks: wild-type N = 6, n = 36, SCA3 N = 8, n = 32. 34-weeks: wild-type N = 12, n = 29, SCA3 N = 16, n = 40. 60-weeks: wild-type N = 5, n = 25, SCA3 N = 5, n = 25. *p<0.05, **p<0.01, ***p<0.001. T-test (F – all. G – 12-weeks. M – all. N – 12-weeks, 60-weeks), Mann-Whitney test (E – all. G – 34-weeks, 60-weeks. L – all. N – 34-weeks).

### Purkinje Cell Activity is Irregular in SCA3 Mice

The abnormal DCN firing can be caused from irregular firing of DCN neurons themselves, and from irregular input from neurons upstream of the output nuclei. Purkinje cells form the sole output of the cerebellar cortex and converge onto DCN. To determine if in SCA3 mice Purkinje cell firing is altered, Purkinje cell activity was recorded *in vivo* in awake, head-fixed, stationary mice. While it would be surprising to have progressive irregularity in the Purkinje cell activity that was not reflected in the DCN activity, the Purkinje cell activity was assessed for progressive neuronal dysfunction at all stages of disease.

At 12-, 34-, and 60-weeks, Purkinje cells in SCA3 mice had an increased ISI CV that was not progressive with age (12-weeks: wild-type 0.392±0.113, SCA3 0.570±0.261, p<0.0001; 34-weeks: wild-type 0.448±0.077, SCA3 0.531±0.183, p=0.0330; 60-weeks: wild-type 0.472±0.080, SCA3 0.646±0.262, p=0.0112), indicating irregularity of firing in SCA3 Purkinje cells compared with that of the wild-type mice (Figure 2L). Similar to the deep cerebellar nuclei, Purkinje cells in 12-week-old SCA3 mice had a decreased average firing rate compared to wild-types, with a trend to decrease at 34- and 60-weeks (12-weeks: wild-type 64.6±19.2, SCA3 54.9±20.5, p=0.0468; 34-weeks: wild-type 71.0±16.3, SCA3 61.9±20.7, p=0.0541; 60-weeks: wild-type 76.3±24.8, SCA3 64.4±21.2, p=0.0741) (Figure 2M). With the observed decrease in average firing rate, this increased ISI CV likely represents a more pronounced irregularity of Purkinje cells. Taken together, this *in vivo* data demonstrated Purkinje cell dysfunction was present at an early age, but not progressive.

### Synaptic and Intrinsic Components Contribute To SCA3 Purkinje Cell Dysfunction

Both the spontaneous intrinsic activity of Purkinje cells and synaptic activity can contribute to the Purkinje cell irregular firing observed *in vivo*. While motor encoding activity was reduced by recordings in standing, stationary animals, synaptic input is still present *in vivo*. In addition, *in vitro* synaptic input can affect Purkinje cell firing through the spontaneous activity of neurons. Sensorimotor integration in the cerebellum occurs via the glutamatergic and GABAergic inputs onto Purkinje cells. Recording Purkinje cell firing *in vitro* from acute cerebellar slices with synaptic transmission intact or blocked with inhibitors of fast GABAergic and glutamatergic transmission began to disentangle whether the Purkinje cell irregularity observed *in vivo* was from synaptic or intrinsic sources.

Acute sagittal cerebellar sections were obtained from 12- and 34-week-old mice and extracellular single-unit recordings were made from visually identified Purkinje cells. Recordings were not made from 60-weeks of age due to technical difficulties maintaining mice to that age. When synaptic transmission was intact, the ISI CV of SCA3 Purkinje cells was significantly increased at both 12- and 34-weeks (12-weeks: wild-type 0.0787±0.0231, SCA3 0.1095±0.0449, p=0.0002. 34-weeks: wild-type 0.0942±0.0239, SCA3 0.1277±0.0439, p<0.0001) (Figure 3D) while the average firing rate did not change (12-weeks: wild-type 40.7±17.7, SCA3 43.1±14.8, p=0.1919. 34-weeks: wild-type 47.3±17.6, SCA3 55.2±26.8, p=0.0743) (Figure 3E). The increased ISI CV was not progressive, as exprected by the *in vivo* data.

**Figure 3:**
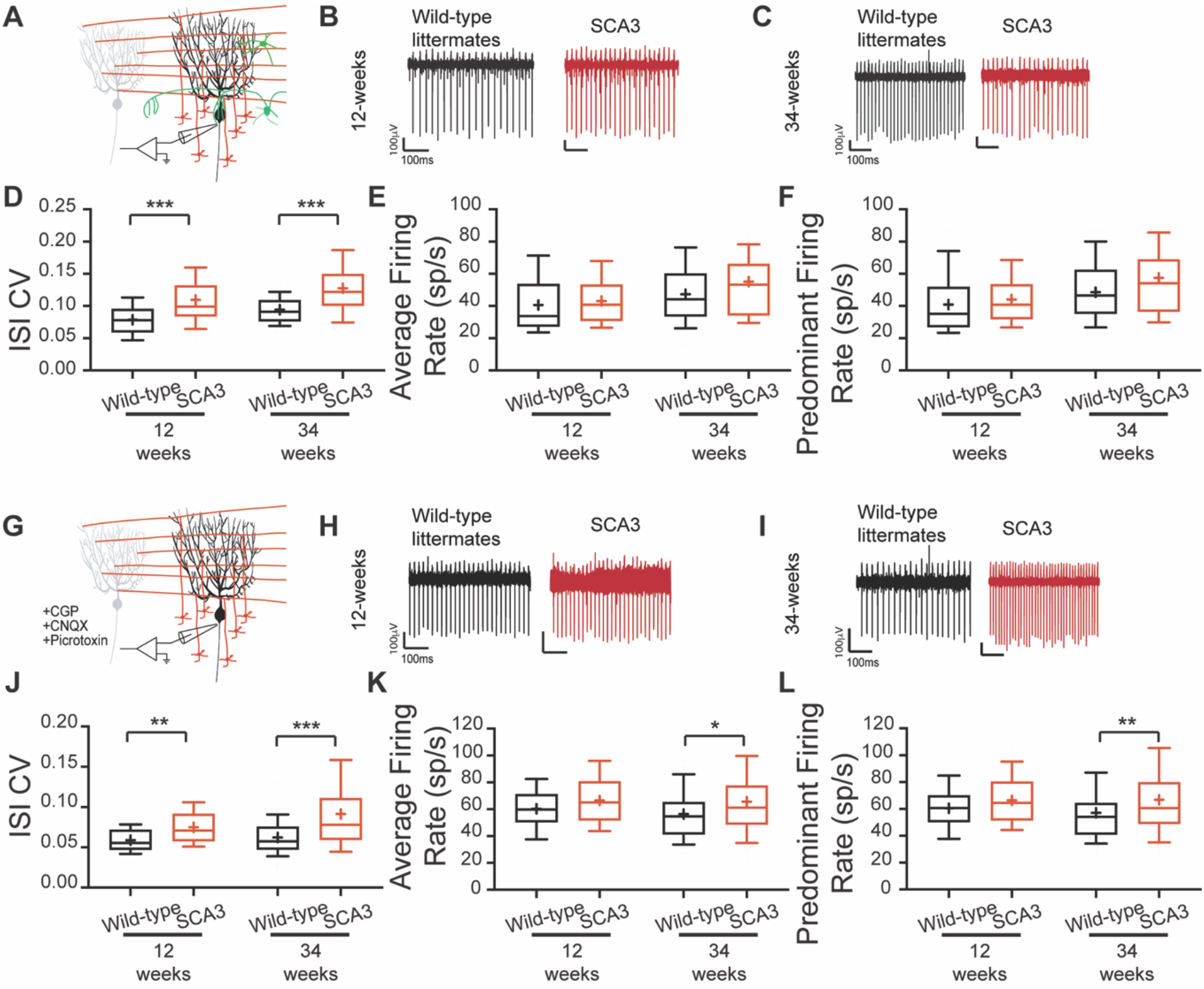
Synaptic And Intrinsic Components To Purkinje Cell Irregularity. *In Vitro* recordings were performed at 12 and 34 weeks of age, with synaptic transmission intact (A-F) and synaptic transmission blocked (G-L). Example Purkinje cell firing traces from 12-weeks (B,H) and 34-weeks (C,I) WT (black) and SCA3 (red). When synaptic transmission is intact (A-F), Purkinje cells ISI CV (C) is increased in SCA3 at both time points with no change in average (E) and predominant (F) firing rate. When synaptic transmission is blocked (G-L), Purkinje cells ISI CV (J) is increased in SCA3 at both time points with no change in average (K) and predominant (L) firing rate at 12-weeks, but an increase in both at 34-weeks. Synaptic transmission intact: 12-weeks: WT N = 4, n = 38; SCA3 N = 4, n = 50. 34-weeks: WT N = 16, n = 89; SCA3 N = 17, n = 91. Synaptic transmission blocked: 12-weeks: WT N = 3, n = 22; SCA3 N = 3, n = 29. 34-weeks: WT N = 13, n = 92; SCA3 N = 15, n = 121. *p<0.05, **p<0.01, ***p<0.001. t-test (K – 12-weeks, L – 12-weeks), Mann-Whitney test (D – all. E – all. J– all. K – 34-weeks. L – 34-weeks).

To determine if intrinsic Purkinje cell dysfunction contributed to the increased ISI CV *in vitro*, fast synaptic transmission was blocked with GABA_A_, GABA_B_, and AMPA/Kainate inhibitors. With synaptic transmission blocked, the ISI CV of Purkinje cells from SCA3 mice remained increased, but was not progressive, at 12- and 34-weeks (12-weeks: wild-type 0.0589±0.016, SCA3 0.0750±0.021, p=0.0040. 34-weeks: wild-type 0.0623±0.022, SCA3 0.0914±0.482, p<0.0001) (Figure 3). The average firing rate of intrinsic Purkinje cell activity had a trend to increase at 12-weeks and was significantly increased at 34-weeks (12-weeks: wild-type 60.3±14.6, SCA3 66.5±19.0, p=0.2057. 34-weeks: wild-type 56.5±20.6, SCA3 65.6±28.0, p=0.0113) (Figure 3D). The increased average firing rate suggests that the firing rate contributed to the increased ISI CV. Therefore, a difference in ISI CV when synaptic transmission was intact suggested that while both synaptic and intrinsic dysfunction occurred, the synaptic component was greater.

## Discussion

Here we show that early in disease the activity of neurons in the DCN of SCA3 mice was irregular with irregular Purkinje cell activity likely a contributing factor. The irregularity in the output of the cerebellum, the DCN, was noted in mice as early as 12-weeks postnatal, however, this irregularity was not progressive with age. Cerebellar Purkinje cells were also irregular *in vivo* as early as 12-weeks, but the irregularity was again not-progressive with age. With *in vitro* experiments, we found that in SCA3 both aberrant intrinsic firing and synaptic activity likely contribute to Purkinje cells dysfunction. The irregular firing in the DCN and Purkinje cells results in a decreased ability to accurately encode motor-related information. Therefore, despite the irregular firing not progressing with motor impairment, the early dysfunction, before any detectable pathology, suggest early, and consistent, cerebellar neuronal dysfunction likely driving the early motor phenotypes in SCA3. With patients, this motor impairment can impact many aspects of their quality of life, as simple things such as dressing and eating require motor coordination. Therefore, with the goal of improving the quality of life of patients, this early Purkinje cell dysfunction could be a good therapeutic target for SCA3 patients.

### Early Cerebellar Dysfunction Contributes to SCA3 Phenotype

While the motor phenotype in SCA3 mice is progressive, the neuronal dysfunction in DCN and Purkinje cells appeared to be consistent from 12-to-60 weeks of age. If an increased cerebellar dysfunction was the primary factor in causing the progressive ataxia in SCA3, one might expect that as the motor symptoms progressed, the neuronal dysfunction, or ISI CV, would also increase and correspond to the increased symptom severity (Tara *et al*., 2018). However, in SCA3, unlike pure cerebellar ataxias, the brainstem, spinal cord, muscles, and other brain regions, such as the substantia nigra, are all affected (Woods & Schaumburg, 1972; Rosenberg *et al*., 1976; Romanul *et al*., 1977; Stefanescu *et al*., 2015; Hernandez-Castillo *et al*., 2018; Koeppen, 2018; Wang *et al*., 2020). Indeed, in this SCA3 mouse model, it is a combination of these factors that contribute to the presentation of motor symptoms. SCA3 mice have motor incoordination as early as 12-weeks of age, and strength-related behavioral phenotypes that were previously shown as early as 6-weeks. By 60-weeks of age, mice are unable to properly support their body weight, often pulling their body to move instead of using proper ambulation (Teixeira-Castro *et al*., 2015). Additional studies will be needed to determine if the dysfunctional cerebellum is driving other sites of dysfunction in the central nervous system, or if these other areas undergo independent dysregulation. Given the ubiquitous expression of ataxin-3 in the nervous system of this mouse model (as in human patients), it might also be that the spinal cord/peripheral nervous system and striatum/substantia nigra related mechanisms of dysfunction arise separately from the cerebellar mechanisms.

Despite the non-progressive nature of cerebellar irregular firing, the early initial dysfunction would be a great therapeutic target. The cerebellum relies on the regular, spontaneous activity of the DCN to accurately encode movement related information through rapid changes, up or down, in the firing rate of the DCN (Eccles, 1967; Thach, 1968). When the spontaneous activity of the cerebellum is no longer precise, the encoding of information is impaired, a feature apparent in other cerebellar ataxias (Walter *et al*., 2006; Alvina & Khodakhah, 2010b; Tara *et al*., 2018). The SCA3 mice have aberrant spontaneous and synaptic dysfunction of Purkinje cells likely contributing to the altered DCN activity at an early stage of disease when motor incoordination is present. Targeting this neuronal activity therapeutically might provide for a consistent, alas partial, reduction in the severity of ataxia throughout the disease. For patients, motor incoordination impacts most aspects of daily life. In fact, as expected, several components of the scale for assessment and rating of ataxia (SARA) depend on coordination, such as gait, finger-to-nose test, fast alternating hand movements, and finger chase (Schmitz-Hubsch *et al*., 2006; Jacobi *et al*., 2020). All of these measurements will affect daily tasks, such as dressing, grooming, and eating. As SCA3 patients have a decreased quality of life early in disease (Bolzan *et al*., 2021), targeting this early dysfunction could alleviate some of these factors potentially improving their quality of life.

### Purkinje Cell Dysfunction as One Potential Mechanism of SCA3 Pathophysiology

In SCA3 patients, Purkinje cells show limited degeneration, if at all (Woods & Schaumburg, 1972; Rosenberg *et al*., 1976; Seidel *et al*., 2012a; Seidel *et al*., 2012b). Here, in a mouse model that does not show nuclear inclusions nor degeneration of Purkinje cells, we found that Purkinje cells have irregular firing since the first stages of disease. One interesting feature of this irregularity was the dependence on location of the Purkinje cells within the cerebellum (Supplemental Figure 1). Purkinje cells are not a homogenous population, with differential gene expression in stripes throughout the cerebellum (Cerminara *et al*., 2015). One differentially expressed gene, zebrin II, is typically used as a guide for this heterogeneity, with lobules I-V being predominantly zebrin II negative, while lobules VI-X are predominantly zebrin II positive (Cerminara *et al*., 2015). With this rough generalization for gene expression, the Purkinje cell data from 34-week-old animals was sorted by the lobule in which it was located. In this preliminary analysis, we found that cells in lobules VI-X had irregular firing, but not those in lobules I-V (Supplemental Figure 1). Further experiments are needed to directly correlate the changes in the firing pattern of Purkinje cells with zebrin II expression, and to delineate the reasons for this difference.

There are a few theories underlying SCA3 pathophysiology, ranging from the toxicity of protein aggregates/intranuclear inclusions to extensive cell death. One of the most consistently affected regions in pathological analysis of SCA3 patients is the DCN (Woods & Schaumburg, 1972; Rosenberg *et al*., 1976; Seidel *et al*., 2012a; Seidel *et al*., 2012b). In the SCA3 mouse model used here, DCN present with intranuclear aggregates after 24-weeks, and DCN volume loss around 42-weeks (Silva-Fernandes *et al*., 2014). At 12-weeks of age, when intranuclear inclusions are not yet visible and there is no cell death, DCN neurons *in vivo* had irregular activity. While the possibility that this irregular activity is partially driven by DCN neurons themselves remains to be examined, similar to some other cerebellar ataxias, Purkinje cells in SCA3 mice have irregular firing. It is interested to note that erratic firing of Purkinje cells can lead to DCN degeneration, as in other mouse models of movement disorders, DCN pathology can be seen at the late stages of disease even when the causal mutation is isolated to Purkinje cells, or largely drives Purkinje cell specific irregularity (Todorov *et al*., 2012; Fremont *et al*., 2015). Together, this suggests the Purkinje cell dysfunction might not only be a contributor to the DCN irregularity but might lead to the DCN pathology observed later in disease. Additional studies are required to understand the mechanisms that cause the aberrant synaptic and intrinsic dysfunction of Purkinje cells. Understanding the contributors to cerebellar dysfunction might inform potential mechanisms driving all neuronal dysfunction in SCA3, which would allow the development of a multi-faceted approach to treating this complex disease.

## Methods

### Animals

Experiments were performed on 12-to-70-week-old CMVMJD135 (SCA3) mice with 140±10 CAG repeats, bred and genotyped at the Life and Health Science Research Institute, School of Medicine, University of Minho, Braga, Portugal; ICVS/3B’s – PT Government Associate Laboratory. Mice were housed at weaning in groups of 5-6 animals in filter-topped polysulfone cages 267 × 207 × 140 mm (370 cm2 floor area) (Tecniplast, Buguggiate, Italy), with corncob bedding (Scobis Due, Mucedola SRL, Settimo Milanese, Italy) in a SPF animal facility. All animals were maintained under standard laboratory conditions: an artificial 12h light/dark cycle (lights on from 8 a.m. to 8 p.m.), with a room temperature of 21±1°C and a relative humidity of 50–60%). Mice were given a standard diet (4RF25 during the gestation and postnatal periods, and 4RF21 after weaning) (Mucedola SRL, Settimo Milanese, Italy) and water ad libitum. Health monitoring was performed according to FELASA guidelines, confirming the Specified Pathogens status of sentinel animals maintained in the same animal room. All procedures were conducted in accordance with European regulations (European Union Directive 86/609/EEC). Animal facilities and the people directly involved in vertebrate animal experiments (S.D.S, A.N.C) as well as coordinating the research (P.M.) were certified by the Portuguese regulatory entity (Direcção Geral de Alimentação e Veterinária). All protocols performed were approved by the Animal Ethics Committee of the Life and Health Sciences Research Institute, University of Minho and by the DGAV (reference 020317).

Mice were shipped to Albert Einstein College of Medicine and allowed at least 2 weeks of recovery before any experiments were conducted. All experiments were conducted in accordance with the guidelines set by the Institute of Animal Safety and Institute of at Albert Einstein College of Medicine under the senior investigator’s (K.K.) Institutional Animal Care and Use Committee approved protocol. Mice were housed on a 12:12 hour reversed light/dark cycle. Data was analyzed to examine any effect of repeat length on the electrophysiology, with no effect detected. As there have been no reported sex differences in these mice (Silva-Fernandes *et al*., 2010; Teixeira-Castro *et al*., 2011; Duarte-Silva *et al*., 2014; Silva-Fernandes *et al*., 2014; Esteves *et al*., 2015; Teixeira-Castro *et al*., 2015; Duarte-Silva *et al*., 2018; Esteves *et al*., 2019), only female mice were used for Figures 1-2, with Figure 3 including both male and female mice, with no difference between sex detected. For the 12-week time point, mice were 12-to-15-weeks old. For the 34-week time point, mice were 34-to-40-weeks old. For the 60-week time point, mice were 60-to-70-weeks old.

### Behavior

All experiments were performed during the mouse’s dark cycle. For all behavior experiments, the experimenter was blinded to the genotype of the mice.

#### Parallel Rod Floor Test

Mice were assessed on the parallel floor rod test for 10-minute sessions, once a day for two days (Kamens *et al*., 2005; Kamens & Crabbe, 2007). The trials were averaged to obtain total distance traveled and number of foot slips per animal. Ataxia ratio is defined as the number of foot slips (errors) divided by distance traveled (cm).

#### Disability Score

Mice were assessed using the previously published disability score as follows: 0 = normal motor behavior; 1 = slightly slowed or abnormal movements; 2 = mild impairments, limited ambulation unless disturbed; 3 = moderate impairment, limited ambulation even when disturbed, frequent abnormal postures; 4 = severe impairment, almost no ambulation, sustained abnormal postures; 5 = prolonged immobility in abnormal postures (Weisz *et al*., 2005; Tara *et al*., 2018). To assess the disability score, mice were individually placed in an open field for 30 minutes. The videos were blinded, and a 30-second segment was selected where the mouse was ambulating. The cropped video was sent to 4 viewers trained on the disability scale and blind to the genotype of the animal. These 4 scores were averaged to produce a score for each trial. The score for each mouse was the average of at least 2 trials from separate days.

### Electrophysiology

#### In vivo

Mice were implanted with a titanium bracket fixed to the skull with optibond (Kerr Dental) and charisma (Kulzer). A recording chamber was formed on the skull above the cerebellum with dental cement. Craniotomies for recordings were made at the following locations (A/P, M/L): −6.2, ±1.75; and −7, ±0.25. Until time of recording, craniotomies were covered with silicone adhesive (KWIK-SIL, WPI). From awake, head-fixed, non-moving mice, extracellular single-unit activity was recorded by advancing a tungsten electrode (Thomas Recordings, 2-3 MΩ) until Purkinje cell layers or deep cerebellar nuclei were detected. Purkinje cells were identified by location, presence of complex spikes, and characteristic firing rate. Deep cerebellar nuclei were identified by location and characteristic firing rate. Signals were filtered (200 Hz-20 kHz) and amplified (2000x) using a custom-built amplifier, and then digitized (20 kHz) using a National Instruments card (PCI-MIO-16-XE) with custom written software in Labview. Waveforms were sorted offline using amplitude, energy, and principal component analysis (Plexon offline sorter).

#### In vitro

All experiments were conducted blinded to the genotype of the mouse. Mice were anesthetized with isoflurane and rapidly decapitated. The brain was rapidly removed and placed in warm (35±2°C) artificial cerebral spinal fluid (ACSF): (in mM) NaCl 125, KCl 2.5, NaHCO_3_ 26, NaH_2_PO_4_ 1.25, MgCl_2_ 1, CaCl_2_ 2, glucose 11, pH 7.4 when gassed with 5% CO_2_:95% O_2_. The cerebellum was dissected and 300 μm thick sagittal sections were made using a Campden instruments 7000sz vibratome. Slices were kept in oxygenated ACSF at 35±2°C for 1 hour and then kept at room temperature until use (up to 4 hours). At the time of recording, slices were placed in a recording chamber perfused with warm (35±2°C) ACSF at 1.5 ml/min. Single-unit extracellular recordings were obtained from visually identified Purkinje cells with an upright microscope (Zeiss) using a home-made differential amplifier and glass pipette back-filled with ACSF. Where indicated, to isolate Purkinje cell intrinsic activity, perfusion ACSF contained 10μM Picrotoxin, 1μM CGP55845, and 10μM CNQX to block GABA_A_, GABA_B_, and AMPA/Kainate receptors respectively. Signals were digitized (10 kHz) using a National Instruments card (PCI-MIO-16-XE) with custom written software in Labview. Waveforms were sorted offline using amplitude, energy, and principal component analysis (Plexon offline sorter), and output was analyzed using custom written Labview software. For all experiments, each cell was held for a minimum of 5 minutes.

### Statistics

Data is graphed using 10-90 box plots, with the mean indicated by the plus sign. T-tests were used for all pair-wise comparisons. All data was assessed for normalcy with the Shapiro-Wilk test, and non-parametric tests (Mann-Whitney) were conducted for all datasets that were not normally distributed. All data reported in the text as ± standard deviation. ‘N’ refers to number of mice. ‘n’ refers to number of cells. * p<0.05, **p<0.01, ***p<0.001.

---

**Supplemental Figure 1:**
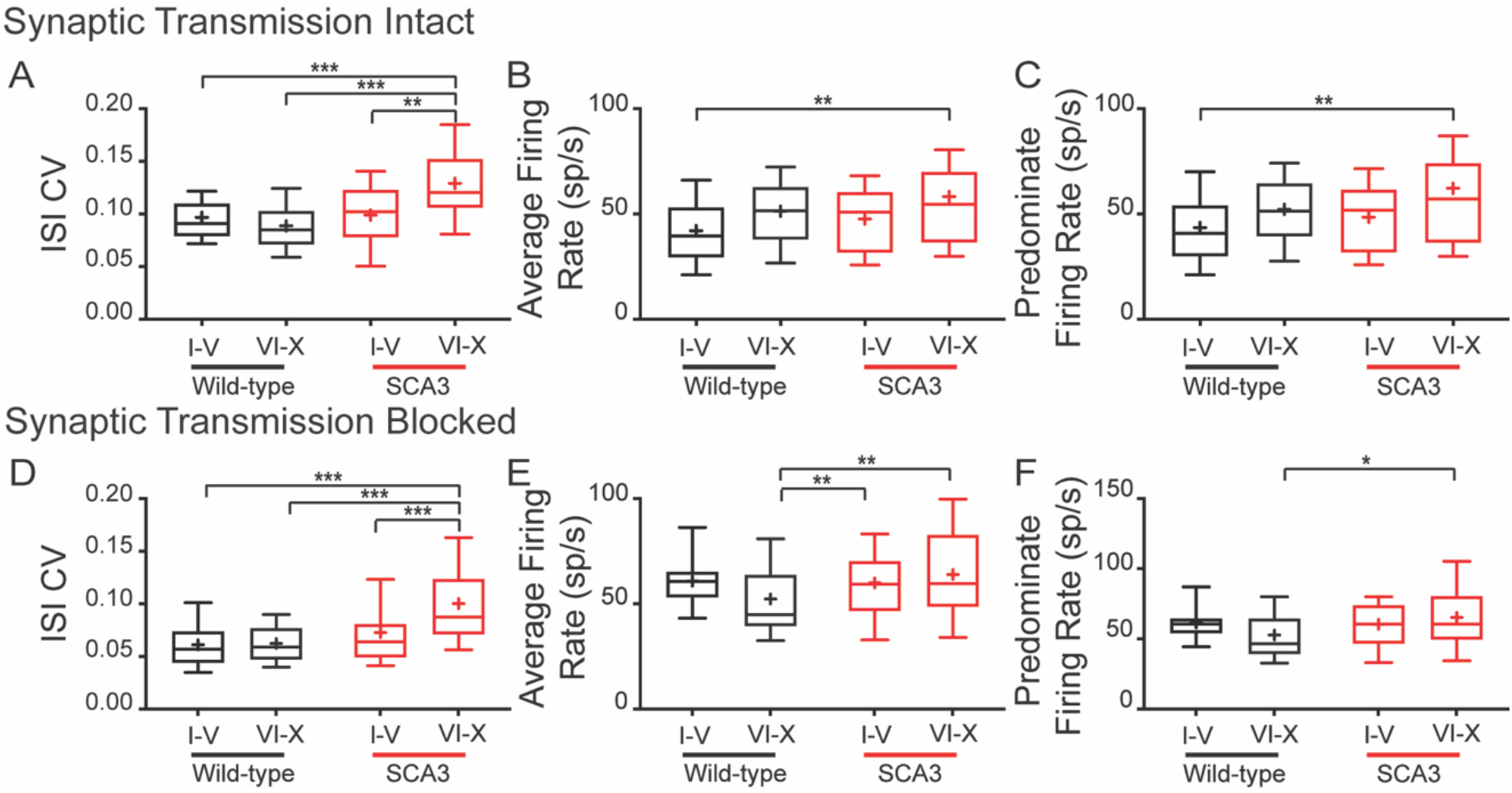
SCA3 Purkinje Cell Dysfunction is Heterogeneous. The *in vitro* Purkinje cell recordings from 34-week-old animals demonstrated dependence on location of recording. When synaptic transmission was intact (A-C), Purkinje cell ISI CV (A) was increased with a small increase in average (B) and predominant (C) firing rates only in SCA3 lobules VI-X. This held true when synaptic transmission is blocked (D-F). The increased ISI CV (D), increased average firing rate (E) and increased predominant firing rate is specific to SCA3 lobules VI-X. Synaptic transmission intact: wild-type I-V: N = 10, n = 43; wild-type VI-X: N = 10, n = 33. SCA3 I-V: N = 5, n = 14; SCA3 VI-X: N = 13, n = 54. Synaptic transmission blocked: wild-type I-V: N = 7, n = 29; wild-type VI-X: N = 12, n = 59. SCA3 I-V: N = 11, n = 43; SCA3 VI-X: N = 14, n = 59. *p<0.05, **p<0.01, ***p<0.001. One-way ANOVA with Bonferroni correction for multiple comparisons.

